# DyGraphTrans: A temporal graph representation learning framework for modeling disease progression from Electronic Health Records

**DOI:** 10.64898/2026.01.28.702347

**Authors:** Most Tahmina Rahman, Mohammad Al Olaimat, Serdar Bozdag, Alzheimer’s Disease Neuroimaging Initiative

**Author notes:** Data used in preparation of this article were obtained from the Alzheimer’s Disease Neuroimaging Initiative (ADNI) database (adni.loni.usc.edu). As such, the investigators within the ADNI contributed to the design and implementation of ADNI and/or provided data but did not participate in the analysis or writing of this report. A complete listing of ADNI investigators can be found at: http://adni.loni.usc.edu/wp-content/uploads/how_to_apply/ADNI_Acknowledgement_List.pdf.

## Abstract

**Motivation:** Electronic Health Records (EHRs) contain vast amounts of longitudinal patient medical history data, making them highly informative for early disease prediction. Numerous computational methods have been developed to leverage EHR data; however, many process multiple patient records simultaneously, resulting in high memory consumption and computational cost. Moreover, these models also often lack interpretability, limiting insight into the factors driving their predictions. Efficiently handling large-scale EHR data while maintaining predictive accuracy and interpretability therefore remains a critical challenge. To address this gap, we propose DyGraphTrans, a dynamic graph representation learning framework that represents patient EHR data as a sequence of temporal graphs. In this representation, nodes correspond to patients, node features encode temporal clinical attributes, and edges capture patient similarity. DyGraphTrans models both local temporal dependencies and long-range global trends, while a sliding-window mechanism reduces memory consumption without sacrificing essential temporal context. Unlike existing dynamic graph models, DyGraphTrans jointly captures patient similarity and temporal evolution in a memory-efficient and interpretable manner.

**Results:** We evaluated DyGraphTrans on Alzheimer’s Disease Neuroimaging Initiative (ADNI) and National Alzheimer’s Coordinating Center (NACC) for disease progression prediction, as well as on the Medical Information Mart for Intensive Care (MIMIC-IV) dataset for early mortality prediction. We further assessed the model on multiple benchmark dynamic graph datasets to evaluate its generalizability. DyGraphTrans achieved strong predictive performance across diverse datasets. We also demonstrated interpretability of DyGraphTrans aligned with known clinical risk factors.

The source code and datasets are available at https://github.com/bozdaglab/DyGraphTrans.

**Supplementary:** Uploaded as an attachment.

## 1 Introduction

The availability of large-scale multimodal electronic health records (EHRs) presents significant opportunities for predicting disease diagnosis and susceptibility. Many deep learning and machine learning methods have been used for disease diagnosis or subtype prediction (Yadav and Jadhav, 2019; Ha et al., 2019). However, these approaches typically rely on a single type of data modality such as imaging data or laboratory tabular data. In practice, most disease conditions can be influenced by many different factors, such as physiological changes, clinical treatments, and evolving health states, making disease prediction a challenging task. Therefore, effectively integrating multiple data modalities is essential for improving disease prediction performance.

Recently, static graph-based models have been used to integrate multimodal datasets. Unlike traditional deep learning approaches, graph methods capture patient–patient similarity by connecting individuals with similar multimodal biomedical data and leverage information from biologically and clinically close neighbors. For instance, Similarity Network Fusion (SNF) (Wang et al., 2014) integrates multiple data modalities such as mRNA expression, DNA methylation, and microRNA expression data. It constructs patient similarity networks for each modality and iteratively fuses them into a combined network representation. This allows SNF to capture complementary information from each dataset. MOGONET (Wang et al., 2021) and SUPREME (Kesimoglu and Bozdag, 2023) leverage graph convolutional networks (GCN) to learn relationships across data types for cancer subtype prediction and achieve high precision. These approaches rely on static datasets, but patient conditions or disease characteristics can change over time, which highlights the importance of considering the dynamism of the datasets.

To learn from similar nodes in a graph and to handle temporal changes, many dynamic graph-based models such as EvolveGCN (Pareja et al., 2020) and ROLAND (You et al., 2022) have been introduced.EvolveGCN combines GCN and RNN to capture the dynamism of graphs and ROLAND modifies the static graph neural networks (GNN) architecture by adding a update module inside GNN to handle dynamic graphs.

Recent models like GraphSSM (Li et al., 2024) and WinGNN (Zhu et al., 2023) offer more scalable alternatives to dynamic graphs. GraphSSM introduces an extension of state space models using Laplacian-regularized memory compression and mixed discretization to model evolving graph structures. WinGNN removes the need for explicit temporal encoders by leveraging a meta-learning framework and sliding-window gradient aggregation for learning across both short- and long-term scales. However, these architectures face limitations in capturing long-range dependencies and may struggle to accurately capture sudden or abrupt changes in graph structure or node attributes, which are common in clinical data. They also offer limited interpretability, which makes their predictions harder to validate in clinical settings.

Beyond improving predictive performance, some studies have emphasized the need for interpretable EHR models that provide clinically meaningful explanations. RETAIN (Choi et al., 2016) employs two separate RNNs to compute dual-level attention, a visit-level attention that identifies which past encounters are most influential, and a variable-level attention that highlights specific clinical features within each visit that contribute to the prediction. TA-RNN (Al Olaimat et al., 2024) extends traditional RNN-based methods by integrating a temporal-attention mechanism that explicitly emphasizes clinically important time points in a patient’s history. Instead of treating each visit equally, TA-RNN learns attention weights that amplify informative events such as changes in cognitive scores or magnetic resonance imaging (MRI) features. However, TA-RNN relies on abstract hidden states that lack the ability to address sudden changes.

Several GNN-based post-hoc explanation methods, such as GNNExplainer (Ying et al., 2019) and PGExplainer (Luo et al., 2020), aim to identify important subgraphs or edges that drive model predictions. However, these methods are primarily developed for static graph settings and do not explicitly address the challenges of interpretability in dynamic graphs. Yao and Joe-Wong (2021) introduce an interpretable dynamic graph clustering framework which quantifies the relative importance of historical connections in evolving graphs. However, studies on interpretable dynamic graph models remain limited, particularly in the context of EHR datasets.

To address the aforementioned limitations, we introduce DyGraphTrans, a dynamic graph representation learning framework designed to model how patients’ health states evolve over time using longitudinal EHR data. Instead of treating longitudinal patient records as isolated sequences, we represent patients as nodes within a graph, which allows the model to learn from both individual attributes and relationships among clinically similar patients. DyGraphTrans combines modules to capture local and global disease progression patterns. Unlike embedding-based update mechanisms used in many dynamic graph models, DyGraphTrans tracks and learns patterns directly from model weight evolution. As a result, it enables efficient training and inference on large dynamic graphs such as Medical Information Mart for Intensive Care (MIMIC-IV) with a compact parameterization. To improve efficiency, we also incorporate a sliding-window mechanism that limits computation to a subset of recent clinical records while preserving essential temporal context. This approach not only improves scalability but also helps the model focus on the most relevant and up-to-date clinical information. In addition, DyGraphTrans extracts feature-level signals within each temporal window that help identify interpretable insights into clinical factors.

In general, the design of DyGraphTrans is well suited for EHR data because disease trajectories involve both evolving patient states and relationships among similar individual patterns. To contextualize existing dynamic graph approaches, Table 1 summarizes key differences among representative dynamic graph models in terms of what is updated over time, their ability to capture global temporal trends, and interpretability. To demonstrate the effectiveness of our framework, we evaluated DyGraphTrans using two AD datasets, namely the Alzheimer’s Disease Neuroimaging Initiative (ADNI) and the National Alzheimer’s Coordinating Center (NACC) to forecast conversion from MCI to AD at the next clinical visit. We also evaluated DyGraphTrans using an Intensive Care Unit (ICU) dataset called the Medical Information Mart for Intensive Care, version IV (MIMIC-IV) where we utilized EHR data from the first 48 hours of admission to the ICU to predict patient mortality at hour 75. To demonstrate the generalizability of DyGraphTrans on other domains, we evaluated it on six benchmark temporal graph datasets, too.

**Table 1.** Comparison of temporal modeling characteristics in dynamic graph models. Update Mech.: update mechanism, Interp.: interpretability, Transf.: Transformer, and Global Trend indicates whether long-term temporal dynamics are explicitly modeled.

| **Model** | **Update Mech.** | **Updated Item** | **Global Trend** | **Interp.** |
| --- | --- | --- | --- | --- |
| EvolveGCN | RNN | GNN weights | No | Low |
| ROLAND | Update module | Embeddings | No | Low |
| WinGNN | Meta-learning | Gradients | Yes | Low |
| GraphSSM | State-space | Latent states | Yes | Low |
| DyGraphTrans | RNN+Transf. | GNN weights | Yes | High |

We summarize the main contributions of our study as follows:

- We propose DyGraphTrans, which updates GNNs with RNN and Transformer-based weight–evolution modules to model temporal changes in patient histories for early prediction of patient clinical outcomes.
- We incorporate a sliding-window strategy that enables DyGraphTrans to efficiently capture both short-term dynamics and long-range temporal trends of patients while substantially reducing memory consumption.
- We conduct an interpretability analysis that identifies which temporal windows are most influential for the prediction and within those windows which specific features contribute most to the model’s decision.
- We evaluated DyGraphTrans on ADNI, NACC, MIMIC-IV, and six benchmark temporal graph datasets, achieving strong and competitive performance compared to state-of-the-art (SOTA) methods in most cases.

## 2 Materials and Methods

DyGraphTrans is a dynamic graph representation learning framework for longitudinal EHR data, where each clinical time point is modeled as a patient–patient similarity graph with multimodal node features. The model processes sequences of graphs using a GNN that combines an RNN to capture changes between consecutive records and a Transformer to learn long-range temporal patterns. A sliding-window mechanism allows DyGraphTrans to focus on the most relevant recent history while maintaining computational efficiency.

**Preliminaries:** We model longitudinal EHR as a sequence of dynamic patient-patient graphs observed across multiple time points. Let *N* denote the number of patients and we associate each patient with a node in the graph. At each time point *t* ∈ {1, …, *T*}, we construct a graph *G*^(*t*)^ = *V, E*^(*t*)^, *X*^(*t*)^, where the node set *V* = {*v*_1_, …, *v*_*N*_ } remains fixed across all time points. Each patient node *v*_*i*_ is associated with multimodal EHR features collected from *M* data modalities (e.g., demographics, cognitive assessments, and imaging). Let 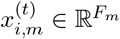 denote the feature vector of patient *i* at time *t* for modality *m* ∈ {1, …, *M* }, where *F*_*m*_ is the dimensionality of modality *m*. The multimodal patient representation is obtained by concatenating modality-specific features: 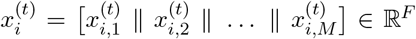, where 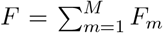 is the total feature dimensionality. Collecting all patient feature vectors yields the node feature matrix *X*^(*t*)^ ∈ ℝ^*N ×F*^. Patient-patient relationships at time *t* are encoded using pairwise similarity between feature vectors. For patients *i* and *j*, we define 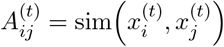, where sim(·, ·) denotes a similarity metric. Here, SNF is used to integrate modality-specific similarity matrices into a single fused adjacency matrix *A*^(*t*)^, from which the edge set *E*^(*t*)^ is derived. Assembling the graphs from all time points gives the temporal graph sequence 𝒢 = {*G*^(1)^, *G*^(2)^, …, *G*^(*T*)^}, which jointly represents the evolution of individual patient states and inter-patient relationships across time points.

**Sliding window:** To efficiently capture temporal dependencies, we employ a sliding window of length *L*, which covers *L* consecutive time points. For example, a window size of *L* = 3 uses time points (1, 2, 3) to predict GNN weights at time 4, the next window uses (2, 3, 4) to predict weights at time 5, and so on.

### 2.1 Datasets

We evaluated our proposed framework on three EHR datasets (i.e., ADNI, NACC, and MIMIC-IV v2.2), four small-scale and two large-scale temporal benchmark graph datasets.

In ADNI, we utilized cross-section demographic, longitudinal cognitive- and MRI-based structured data (https://adni.loni.usc.edu). ADNI was launched in 2003 as a public–private partnership led by Principal Investigator Michael W. Weiner, MD, with the primary goal of determining whether serial MRI, positron emission tomography (PET), biological markers, and neuropsychological assessments can be combined to track the progression of MCI and early AD. Following the preprocessing pipeline described in PPAD (Al Olaimat et al., 2023), we retained cognitive and MRI data from visits at months 0, 6, 12, 24, 36, 48, and 60. Patients with any missing visits were removed, resulting in a final dataset of 137 unique participants.

The second dataset, NACC (Besser et al., 2018), is designed to support research on AD and related dementias by combining data from multiple study sites across the United States. It provides both longitudinal and cross-sectional data, including demographic information, cognitive assessments, medical history, genetic data, and neuroimaging measures. Using the same PPAD preprocessing pipeline, we extracted demographic and cognitive features across eight consecutive visits for 440 unique participants.

For both ADNI and NACC, we used prior visit data to predict MCI-to-AD conversion at the next visit. We used clinical data (without using diagnosis labels) from visits 1–6 to predict visit 7 in ADNI, and visits 1–7 to predict visit 8 in NACC.

Our third cohort was derived from the temporal MIMIC-IV v2.2 database, which provides detailed longitudinal clinical data for respiratory support and ICU management. Specifically, we used the semi-processed version released through PhysioNet (Moukheiber et al., 2025; Moukheiber et al.; Goldberger et al., 2000). This dataset includes 90-day hourly measurements of ventilation status, vital signs, laboratory values, and major treatment interventions, along with outcome variables such as ICU length of stay and mortality for 50,920 patients. Based on average ICU length of stay, we retained the time steps for the first 75 hours for each patient. Features with ≥ 70% missing rate were removed, and patients with more than 40% missing data were excluded. Missing temporal values were imputed using linear interpolation within patients followed by K-nearest neighbors (KNN) imputation across patients. Categorical harmonization included recoding all marital status missing values as “Unknown” and collapsing 33 race categories into five major groups. To reduce class imbalance, we restricted the cohort to patients who received any respiratory intervention such as invasive ventilation, non-invasive ventilation, high-flow oxygen or who had a documented death outcome at any time point. The final processed dataset had 19,594 patients including 2,269 death and 17,325 survived cases. We used patient attributes and relationships of the first 48-hour time points to predict mortality in the hospital at hour 75 without incorporating labels at intermediate time points (see Supplementary Document Section 1.2).

We also evaluated our method four small-scale (DBLP-3, Brain, Reddit, and DBLP-10) and two large-scale (arXiv and Tmall) benchmark datasets. The processed temporal graphs were obtained by directly running the data preprocessing code released with GraphSSM, without additional modifications (Li et al., 2024) (see Supplementary Document Section 1.2).

#### 2.1.1 Building a Homogeneous Patient–Patient Network

For ADNI, three modality-specific networks were generated from demographic, cognitive, and MRI features; for NACC, two networks were built from demographic and cognitive data; and for MIMIC-IV, three networks were constructed based on static demographic variables, laboratory measurements, and combined vital-sign and related laboratory information. Figure 1A illustrates the graph construction. For each time point *t* and each modality *m* ∈ {1, …, *M* }, we first constructed a modality-specific adjacency matrix 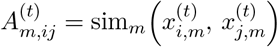 which defines the patient-patient similarity graph for. We utilized the SNF method that iteratively aligns modality-specific similarity graphs to produce a fused network capturing shared structure across data types. It repeatedly exchanges neighborhood information across modalities until convergence. In SNF, the *similarity function* specifies how pairwise relationships are computed such as Jaccard similarity for binary or categorical data (e.g., demographics) and Euclidean distance for continuous features (e.g., cognitive scores, MRI measurements, laboratory values). Each similarity matrix 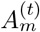 is then converted into an affinity matrix 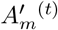. SNF combines these modality-specific graphs into a single fused adjacency: 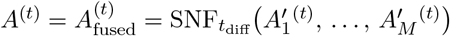. The final homogeneous patient-patient graph at time point *t* is:

**Figure 1.**
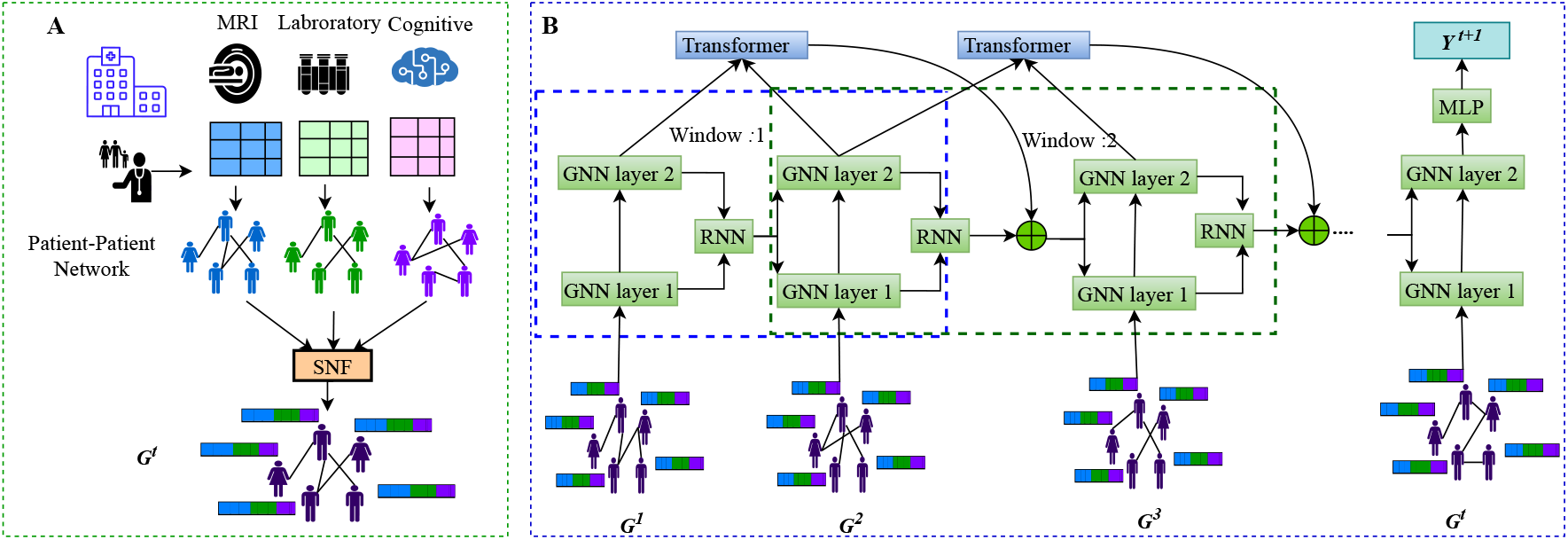
Overview of our proposed DyGraphTrans Model. **A.** Each patient time point is represented as a graph where nodes correspond to patients and edges represent patient similarity computed using SNF on multimodal EHR data. **B**. After message aggregation from neighbors using GNN, RNN module updates the GNN layers weights with previous hidden states. Sequences of GNN weights are processed through a sliding-window mechanism via Transformer. After the first window, GNN weights are updated at every time point with weights generated by RNN and Transformer. Finally, processing all time points till the time point *t*, one classifier, MLP is used for predicting labels *Y* at time point *t+1*.

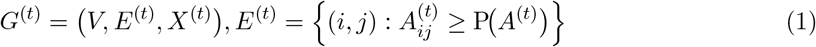

In Eq. 1, *P* (·) represents the percentile similarity threshold. The patient similarity graphs were constructed once during preprocessing and then fixed for all experiments. The dynamic patient graphs were used for DyGraphTrans and all baseline models ensuring that comparisons reflect differences in model learning rather than graph construction. SNF hyperparameter values are provided in Supplementary Table S1. Summary statistics of the final EHR datasets and benchmark dynamic datasets are shown in Supplementary Table S2 and Supplementary Table S3.

### 2.2 DyGraphTrans

DyGraphTrans is designed to classify patients from longitudinal EHR, represented as a temporal sequence of patient-patient graphs. Unlike static GNNs, DyGraphTrans updates its graph convolutional weights at every time point to model how message passing evolves over time. Instead of updating node embeddings, DyGraphTrans models temporal changes directly in the GNN parameters. This choice reduces dependence on graph size, improves scalability for large graphs, and captures how message passing adapts as patient populations and relationships change. Temporal weight evolution is modeled using both an RNN and a FlashAttention Transformer (Dao et al., 2022). The predicted weights are then inserted into the GNN layers for the current time point. The overview of our model is illustrated in Figure 1B.

#### 2.2.1 Graph Feature Encoding

For each time point *t*, DyGraphTrans processes the patient graph *G*^(*t*)^ to extract a meaningful representation that reflects both individual patient attributes and relationships among similar patients. Let *X*^(*t*)^ ∈ ℝ^*N ×F*^ denote the patient attribute matrix at time point *t*, where *N* is the number of patients and *F* is the number of clinical features per patient. A learnable projection matrix *W*_in_ ∈ ℝ^*F ×d*^ maps these raw features into a *d*-dimensional hidden space, producing the initial node representation 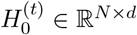. The transformation is computed using a linear projection followed by a ReLU activation: 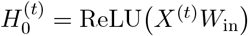. This step maps the raw clinical features into a latent space where nonlinear patterns can be more easily captured. The projection prepares the node features for graph message passing. After that, a stack of GNN layers is applied to propagate clinical information across patients: 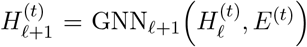. Each GNN layer *ℓ* aggregates information from neighboring patients in the graph. Stacking layers lets the model gather information from farther away. For instance, a 2-layer GNN can gather information from 2-hop neighborhood.

##### Extraction of temporal weight history

To capture how the GNN message-passing behavior GNN evolves over time, we track the changes in its learnable parameters across time points. Let 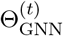 denote the set of learnable GNN parameters at time point *t*. To flatten these parameters into a vector, we apply the vectorization operator vec(.) and obtain 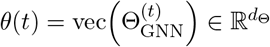 where *d*_Θ_ is the total number of GNN parameters. We construct a sliding window at time point *t*, where each window contains *L* consecutive time points: Ƶ ^(*t*)^ = *θ*(*t* − 1), *θ*(*t* − 2), …, *θ*(*t* − *L* + 1) which stores the GNN parameters in the last *L* time points.

#### 2.2.2 Short-Range Modeling

To effectively model short-term temporal changes and quickly adapt to recent variations in patient condition, DyGraphTrans employs a RNN module that relies only on the previous hidden state and the immediately preceding GNN weight vector to generate the current weight vector. At each time point *t*, the model feeds the most recent GNN weight vector *θ*(*t* − 1) into a recurrent module, which may be an RNN, Gated Recurrent Unit (GRU), or LSTM. Let ℛ denote this generic recurrent function. The recurrent update is written as *h*^(*t*)^, *S*^(*t*)^ = ℛ*θ*(*t* − 1), *S*^(*t*−1)^ where, *θ*(*t*) is the current flattened GNN weight vector, *S*^(*t*−1)^ denotes the previous recurrent state, *h*^(*t*)^ is the output hidden representation produced at time point *t*. The hidden output *h*^(*t*)^ ∈ ℝ^*d*^ is passed through a linear projection 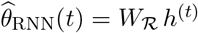 where 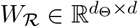 is a trainable matrix. This produces the recurrent module’s estimate of the GNN weights 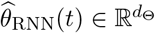 at time point *t*.

#### 2.2.3 Long-Range Modeling

To model long-range temporal dependencies in the evolution of GNN parameters and capture the most critical window for early prediction, we deploy FlashAttention Transformer (Dao et al., 2022) in one window to obtain a window representation. FlashAttention is employed here because it computes self-attention in a memory-efficient and numerically stable manner which provides faster training and scalability to longer temporal windows compared to standard Transformer. If the weight history contains at least *L* entries, the stacked matrix 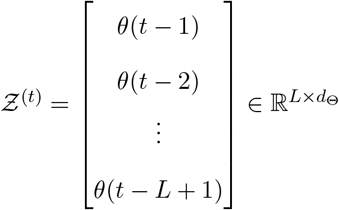 is projected into a hidden space and passed through *B* Transformer encoder layers, each using FlashAttention mechanism. Specifically, we apply a linear projection to map the weight-history matrix into a *d*-dimensional hidden space: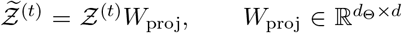 This sequences serve as the input to the Transformer.

Each Transformer layer *b* computes attention using the scaled dot product. A single linear projection produces queries, keys, and values: 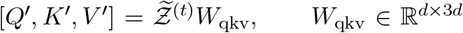 These projections are reshaped into multi-head form: 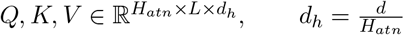, where *H*_*atn*_ is the number of heads and *L* is the sequence length. The scaled dot-product attention scores are: 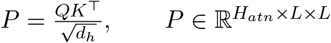. FlashAttention computes the numerically stable softmax in a fused kernel, producing the attention weights *a*(*t*) = softmax(*P*) where 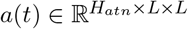. The attention output is: 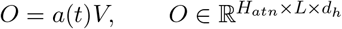, which is concatenated across heads and passed through the output projection to form the layer output 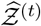. The final representation 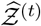 is then mapped to a weight vector 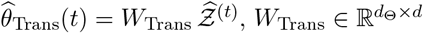, which represents the Transformer module’s estimate of the GNN weights 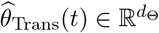 at time point *t*.

#### 2.2.4 Adaptive Fusion of Weight Predictions

To combine the weight dynamics predicted by the recurrent module and the Transformer, the model learns two scalar coefficients, *α* and *β*. These are normalized through a softmax function,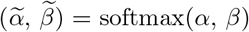 ensuring that 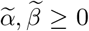 and 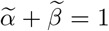. The final predicted weight vector at time *t* is then obtained as a combination of the two sources: 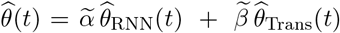 where 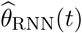 denotes the recurrent module’s GNN weights and 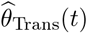 denotes the Transformer’s weights. We replace the GNN’s weights with the weights 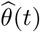 for the current time point, so the GNN updates change over time to reflect evolving patient dynamics.

#### 2.2.5 Prediction Task

After *ℓ* GNN layers at the final time point, the node embeddings 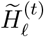 are passed through a linear classifier: 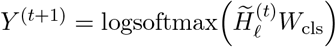. Training loss is applied only at the last time point for EHR data. For benchmark datasets, our model aggregates discrete time level classifier outputs using either the final time point, a weighted average, an exponentially weighted sum, or a uniform mean to produce the final prediction (See Supplemental Document section 1.4).

### 2.3 Attention and Gradient Fusion

To provide clinically meaningful explanations over time for longitudinal EHR, we combine the Transformer’s temporal attention scores with gradient-based feature scores. At each time point *t*, once at least *L* GNN weight vectors are available, the Transformer produces an attention tensor 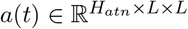 from its final encoder layer. The attention weights are first averaged across heads, and the attention vector corresponding to the final query position is extracted yielding a vector of normalized attention weights 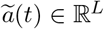 over the *L* time points in the window:

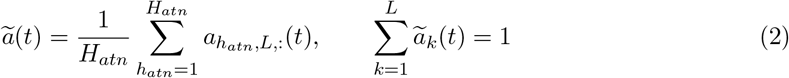

To quantify feature relevance, the supervised loss at the final time point is backpropagated to the node features at each time point, producing gradient magnitudes 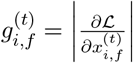. Gradients are summed across nodes to obtain a feature-level vector 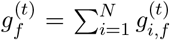. The attention-weighted feature attribution at time *t* is then computed by aggregating feature vectors across the Transformer window using the attention distribution:

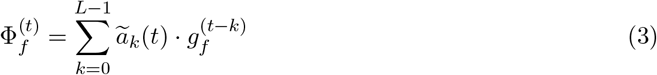

The model provides feature-level temporal interpretability by combining Transformer attention with gradient scores.

## 3 Results and Discussion

Our proposed model, DyGraphTrans, is designed to predict future outcomes from dynamic graphs by capturing local temporal dependencies and long-range global trends within a sliding window. We evaluated our model on three EHR datasets and six benchmark graph datasets. We conducted an ablation study to assess the contribution of each component such as the RNN module, the

Transformer module, and the sliding-window approach. We also demonstrated the interpretability of the model by examining window-level and feature-level scores across the entire test set. Finally, we analyzed the scalability of our model in terms of the number of trainable parameters. The details of these analyzes are described in the following sections.

### 3.1 DyGraphTrans Outperformed SOTA Methods in Predicting Clinical Outcomes

We compared DyGraphTrans with several state-of-the-art dynamic graph- and RNN-based models in three clinical prediction tasks. For all graph-based methods, we used the same series of temporal networks and node features, while for the RNN-based method TA-RNN, we used the patient features only. All models were trained, validated, and tested using identical splits of the dataset and the same random seed to ensure a fair comparison. Due to space limitations, the detailed experimental settings are given in the Supplemental Document Section 1.3 & 1.4.

The results are shown in Table 2. In the ADNI dataset, DyGraphTrans achieved the highest performance surpassing all baselines including the EHR-based TA-RNN and graph-based models such as WinGNN and GraphSSM. On the NACC dataset, DyGraphTrans also outperformed all models in both Micro-F1 and Macro-F1 while methods such as ROLAND and WinGNN perform competitively. For the MIMIC-IV mortality prediction task, which is more acute and highly imbalanced, DyGraphTrans again delivers the strongest results, achieving the best Macro-F1 (0.893) and the highest Micro-F1 (0.960). EvolveGCN-H and TA-RNN showed substantial drops in Macro-F1, which highlighted their difficulty in distinguishing minority severe outcome cases. In contrast, DyGraphTrans maintains high predictive accuracy while effectively capturing short-term changes in ICU settings.

**Table 2.** Performance comparison on ADNI, NACC, and MIMIC-IV datasets (Micro-F1 and Macro-F1, mean *±* std of five runs). Some methods report zero standard deviation because repeated independent runs with the same seed yielded identical performance. Best scores are in **bold**; second-best are underlined.

| Model | ADNI |  | NACC |  | MIMIC-IV |  |
| --- | --- | --- | --- | --- | --- | --- |
|  | Micro-F1 | Macro-F1 | Micro-F1 | Macro-F1 | Micro-F1 | Macro-F1 |
| GraphSSM | 0.921 ± 0.014 | 0.907 ± 0.021 | 0.869 ± 0.022 | 0.845 ± 0.022 | 0.952 ± 0.023 | 0.885 ± 0.055 |
| EvolveGCN-O | 0.964 ± 0.000 | 0.960 ± 0.000 | 0.877 ± 0.018 | 0.844 ± 0.017 | 0.923 ± 0.003 | 0.734 ± 0.019 |
| EvolveGCN-H | 0.950 ± 0.028 | 0.942 ± 0.034 | 0.888 ± 0.004 | 0.855 ± 0.005 | 0.886 ± 0.000 | 0.470 ± 0.000 |
| TA-RNN | 0.928 ± 0.039 | 0.925 ± 0.041 | 0.834 ± 0.065 | 0.723 ± 0.165 | 0.791 ± 0.109 | 0.473 ± 0.004 |
| ROLAND | 0.957 ± 0.014 | 0.952 ± 0.015 | <u>0.922 ± 0.024</u> | <u>0.898 ± 0.034</u> | 0.946 ± 0.012 | 0.850 ± 0.044 |
| WinGNN | <u>0.964 ± 0.000</u> | <u>0.960 ± 0.000</u> | 0.900 ± 0.015 | 0.873 ± 0.017 | 0.949 ± 0.001 | 0.866 ± 0.007 |
| **DyGraphTrans** | **0.971 ± 0.014** | **0.968 ± 0.015** | **0.931 ± 0.007** | **0.913 ± 0.009** | **0.960 ± 0.001** | **0.893 ± 0.003** |

### 3.2 DyGraphTrans Achieved Competitive Performance Across Benchmark Dynamic Graph Datasets

To evaluate the generalizability of DyGraphTrans beyond clinical EHR settings, we further tested the model on six benchmark dynamic graph datasets, including four small-scale (i.e., DBLP-3, Brain, Reddit, and DBLP-10) and two large-scale (i.e., arXiv and Tmall) datasets. For all benchmark experiments, the results of competing methods were taken directly from the GraphSSM paper, and we reproduced our results following the same experimental setup, preprocessing, and evaluation protocol used in their study to ensure a fair comparison (see Supplemental Document Section 1.3 & 1.4). Table 3 summarizes performance in the small-scale dynamic graph datasets. DyGraphTrans achieves the best performance on DBLP-3, DBLP-10 with notable gains over GraphSSM. On the Brain dataset, our method achieves the highest Macro-F1 score while ranking second in terms of Micro-F1. On Reddit, which contains highly dynamic temporal activity, our model remains competitive achieving second-best performance. Performance in large-scale datasets is shown in Table 4. DyGraphTrans achieves the second-best Macro-F1 score on the arXiv dataset. On Tmall, DyGraphTrans delivers competitive performance while remaining stable and memory-efficient. Although GraphSSM attains slightly higher accuracy on Tmall and arXiv by maintaining a latent state for large-scale graphs, this comes at a substantially higher memory cost, limiting scalability.

**Table 3.**
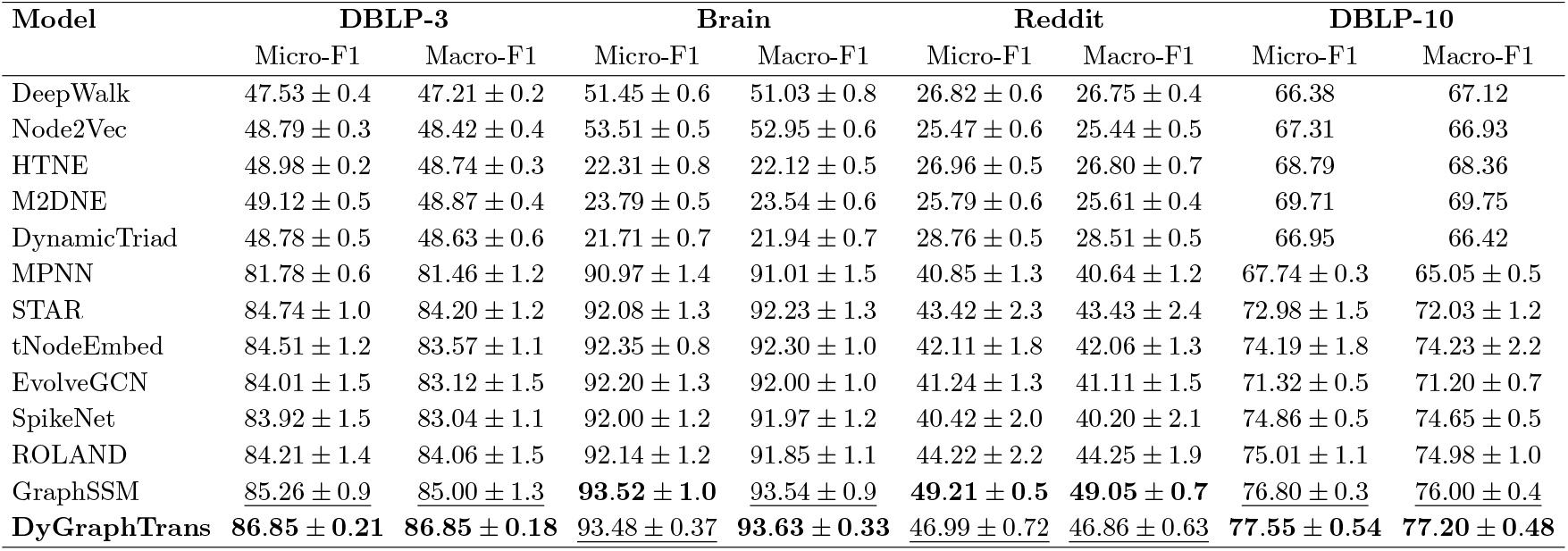
Performance comparison on small-scale dynamic graph datasets. Performance of the baselines are reported in the GraphSSM paper (Li et al., 2024). However, for some baseline methods, standard deviations were not available. Our experiments ensure the identical experimental setting for fair comparisons (Micro-F1 and Macro-F1, mean *±* std of five runs). Best scores are in **bold**; second-best are underlined.

**Table 4.** Performance comparison on large-scale dynamic graph datasets. Performance of the baselines are reported in the GraphSSM paper(Li et al., 2024). Our experiments ensure the identical experimental setting for fair comparisons (Micro-F1 and Macro-F1, mean of five runs). OOM indicates out-of-memory. Best scores are in **bold**; second-best are underlined.

| Model | arXiv |  | Tmall |  |
| --- | --- | --- | --- | --- |
|  | Micro-F1 | Macro-F1 | Micro-F1 | Macro-F1 |
| DeepWalk | 66.54 | 43.01 | 57.88 | 49.53 |
| Node2Vec | 67.71 | 43.69 | 60.66 | 54.58 |
| HTNE | 65.48 | 42.25 | 62.64 | 54.93 |
| M2DNE | 66.91 | 43.52 | 64.65 | 58.47 |
| DynamicTriad | 61.10 | 38.25 | 60.72 | 51.16 |
| MPNN | 64.68 | 41.22 | 58.07 | 50.27 |
| STAR | OOM | OOM | OOM | OOM |
| tNodeEmbed | OOM | OOM | OOM | OOM |
| EvolveGCN | 65.17 | 43.01 | 61.77 | 55.78 |
| SpikeNet | 66.69 | 43.96 | <u>66.10</u> | <u>62.40</u> |
| ROLAND | <u>68.27</u> | 48.01 | OOM | OOM |
| GraphSSM | **70.67** | **49.97** | **66.29** | **62.57** |
| **DyGraphTrans** | 67.28 | <u>48.05</u> | 64.04 | 58.94 |

### 3.3 Ablation Study

To evaluate the impact of each component of DyGraphTrans described in Section 2.2, we performed an ablation study across ADNI, NACC, and MIMIC-IV (Table 5). Removing the RNN results in a model in which the GNN weights are updated solely by the Transformer with a window size of one. In contrast, removing the Transformer yields a model that relies exclusively on the RNN to propagate information over time, without temporal self-attention within the window. Finally, removing the sliding window disables temporal self-attention across multiple time points; in this setting, the Transformer and RNN are applied independently at each time point to update the GNN weights. All experiments were run with identical data splits, fixed random seeds and separately tuned hyperparameters for each ablation setting. Table 5 shows that removing the RNN which models short-term temporal dependencies reduces the performance on NACC where rapid changes dominate early conversion. Excluding the Transformer leads to the largest drops in ADNI confirming its role in capturing long-range disease progression patterns. Removing the sliding window degrades performance in all cases, particularly on MIMIC-IV. These results validate the necessity of each module to achieve the high performance of DyGraphTrans.

**Table 5.** Ablation study results across ADNI, MIMIC-IV, and NACC datasets. Values represent mean *±* standard deviation across five runs. Best scores are in **bold**; second-best are underlined. Transf. denotes Transformer.

| Dataset | Configuration | Micro-F1 | Macro-F1 |
| --- | --- | --- | --- |
| **ADNI** | Without RNN | <u>0.957 ± 0.014</u> | <u>0.952 ± 0.015</u> |
| | Without Transf. | 0.950 ± 0.028 | 0.944 ± 0.031 |
| | Without Window | <u>0.957 ± 0.014</u> | 0.951 ± 0.016 |
|  | DyGraphTrans | **0.971 ± 0.014** | **0.968 ± 0.015** |
| **NACC** | Without RNN | 0.902 ± 0.015 | 0.877 ± 0.017 |
|  | Without Transf. | <u>0.922 ± 0.008</u> | <u>0.903 ± 0.011</u> |
| | Without Window | 0.922 ± 0.023 | 0.901 ± 0.030 |
|  | DyGraphTrans | **0.931 ± 0.007** | **0.913 ± 0.009** |
| **MIMIC-IV** | Without RNN | <u>0.957 ± 0.001</u> | <u>0.888 ± 0.002</u> |
| | Without Transf. | <u>0.957 ± 0.001</u> | 0.887 ± 0.002 |
| | Without Window | 0.944 ± 0.002 | 0.807 ± 0.168 |
|  | DyGraphTrans | **0.960 ± 0.001** | **0.893 ± 0.003** |

### 3.4 Interpretation of DyGraphTrans Models

To assess interpretability, we computed feature and time point scores using our attention–gradient method (see Section 2.3) and aggregated results across 20 independent runs using different seeds. For each feature and temporal window, we report the mean importance weights. The resulting heatmaps for ADNI, NACC, and MIMIC-IV reflect clinically meaningful temporal patterns.

For ADNI, the model assigns the highest importance score to the most recent time point for global cognitive assessments such as CDRSB (Clinical Dementia Rating Sum of Boxes) and FAQ (Total Score of Functional Activities Questionnaire) when predicting MCI to AD conversion (Figure 2). This suggests that recent cognitive status plays a dominant role in the prediction. Additionally, CDRSB receives notable attention even at the earlier time points, which aligns with clinical practice as CDRSB is one of the key measures used by clinicians to diagnose AD (Williams et al., 2013). In contrast, static features such as PTGENDER (gender), PTETHCAT (ethnicity) and PTRACEAT (race) receive relatively low attention across time points. We observed the same trends on the interpretation of the model on the NACC dataset (Supplementary Figure S1). Similar to ADNI, DyGraphTrans consistently assigns high attention to CDRSB across time points and assigns higher attention to the most recent visit compared to earlier visits.

**Figure 2.**
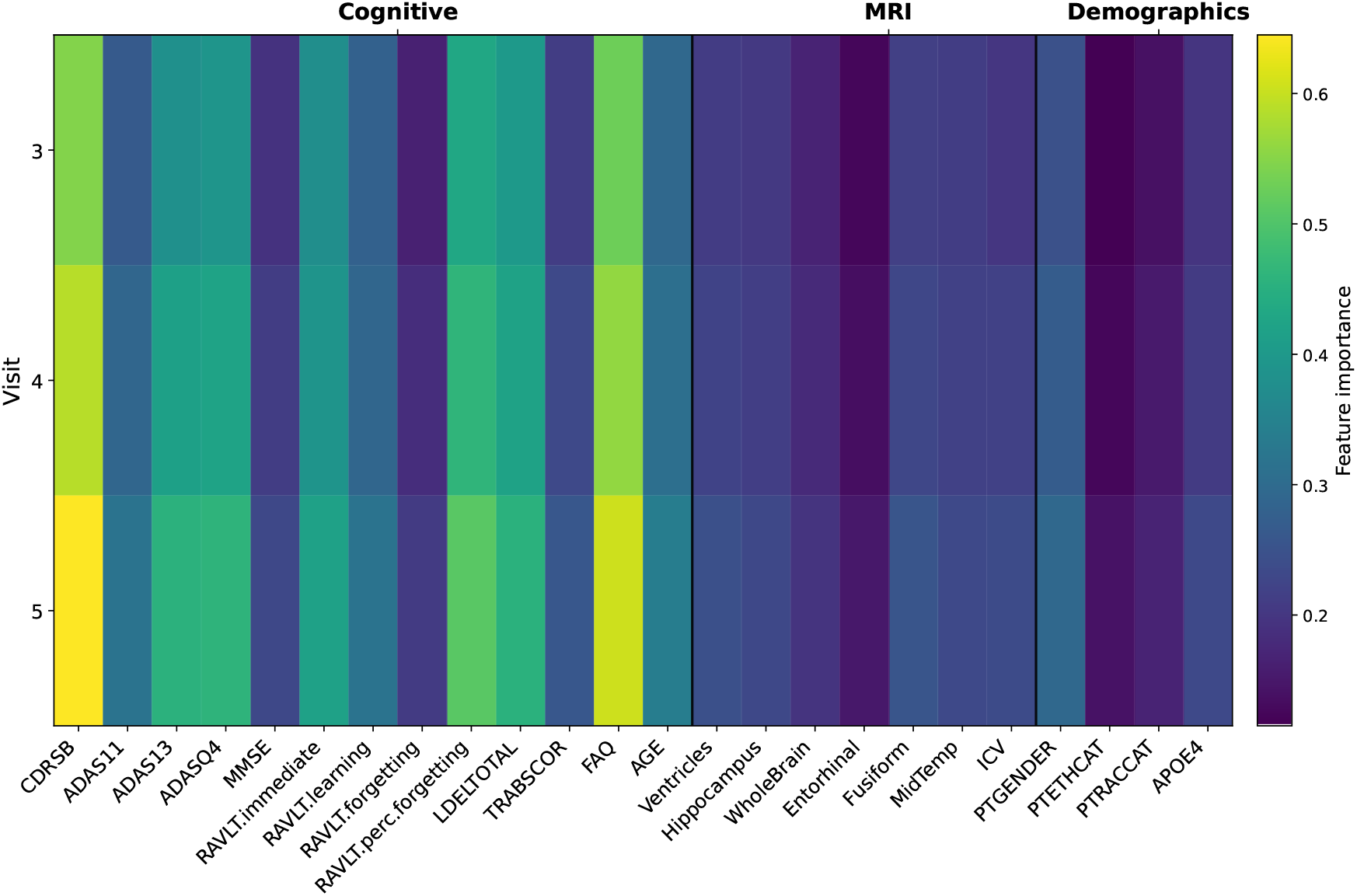
Attention–gradient fused feature importance heatmap for ADNI, aggregated over 20 random seeds (mean values). Each row corresponds to a time point (i.e., clinical visit), and each column corresponds to an EHR feature.

For MIMIC-IV, the model captures the rapidly evolving physiology of ICU patients. In particular, the most informative time points emphasize vital and respiratory features such as ppeak (Peak Inspiratory pressure (cmH2O)), invasive (Invasive ventilation indicator), set fio2 (Set fraction of inspired oxygen), and sofa_24hours (24-hour Sequential Organ Failure Assessment (SOFA) score) (Figure 3), which are well-known indicators of respiratory failure and organ dysfunction in critically ill patients (Moreno et al., 1999; Hung et al., 2018). These features align with known vital markers of deterioration and mortality in the ICU. Additionally, recent and earlier time points receive relatively greater attention, which reflects the importance of initial physiological status at ICU admission as well as recent changes. We also observed that static features receive very low attention scores across the time points showing their limited relevance to patient mortality.

**Figure 3.**
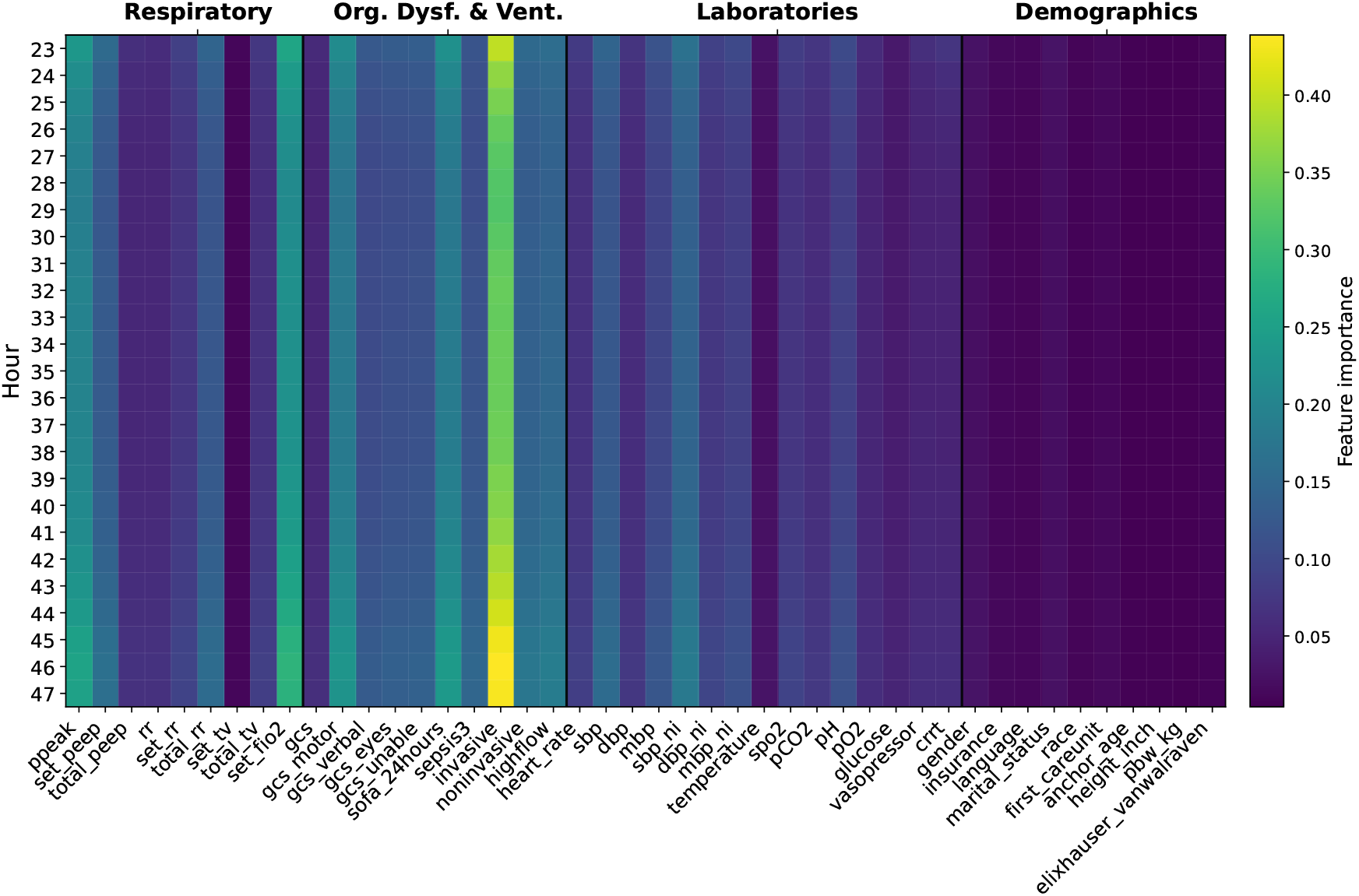
Attention–gradient fused feature importance heatmap for MIMIC-IV, aggregated over 20 random seeds (mean values). Each row corresponds to a time point (i.e., ICU hour), and each column corresponds to an EHR feature. Org. Dysf. & Vent. indicates Organ Dysfunction and Ventilation.

Overall, DyGraphTrans assigns higher importance to known temporal features and consistently low importance weights to demographic and static features, which is expected for disease progression and mortality prediction tasks.

### 3.5 DyGraphTrans Achieved Competitive Performance with Fewer Parameters

In addition to predictive performance and interpretability, we evaluated the scalability of DyGraphTrans by comparing its parameter size against several baselines trained on the ADNI, the NACC, and the MIMIC-IV datasets. For each dataset, every model was independently tuned to achieve its best validation performance before reporting its parameter size. As illustrated in Figure 4, dynamic GNN- and RNN-based baseline models vary widely in their architectural complexity. Several baselines rely on large temporal encoders or recurrent structures that resulted in substantial parameter inflation, such as TA-RNN, which reaches over 700K parameters when trained on the MIMIC-IV. The models such as GraphSSM and ROLAND also were burdened with sizable overhead due to their recurrent or state-space updates. In contrast, DyGraphTrans maintains a compact parameter size across all datasets: 0.9K on the NACC, 3K on the ADNI, and 6K on the MIMIC-IV dataset. Despite incorporating both short-term and long-range temporal modeling, DyGraphTrans remains significantly smaller. This efficiency also came from utilizing FlashAttention Transformer, its sliding-window design, which restricts temporal computation to a limited sequence length, and also from learning temporal patterns in GNN weight trajectories rather than repeatedly encoding full graphs at each time step. Finally, DyGraphTrans achieves a favorable balance of accuracy and scalability while using fewer parameters than many existing dynamic GNN and RNN-based models.

**Figure 4.**
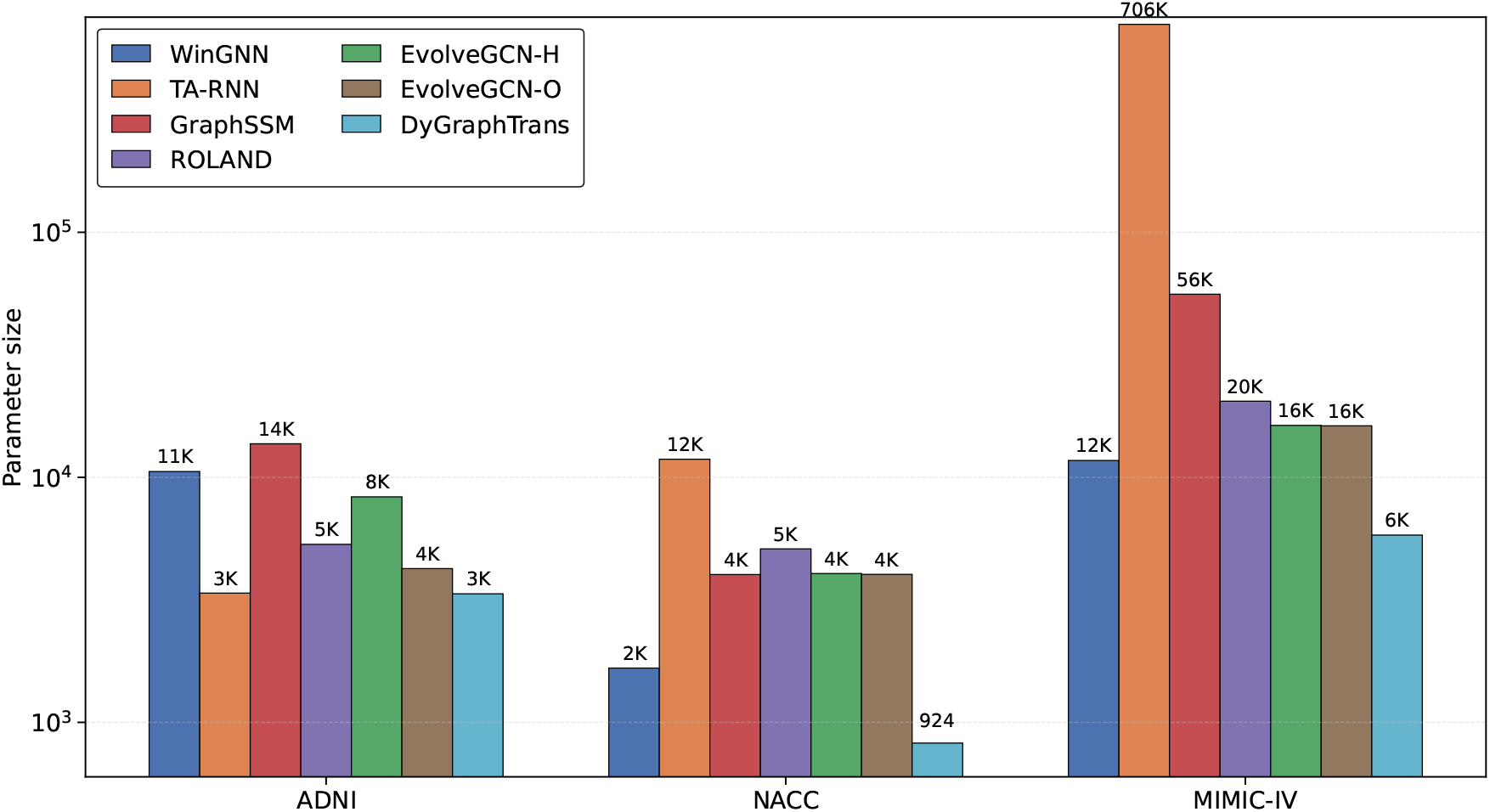
Model parameter sizes across datasets.

## 4 Conclusion

DyGraphTrans efficiently models temporal graphs by integrating GNNs for neighborhood aggregation, RNNs for capturing short-term weight dependencies, and Transformers for learning long-term weight patterns, making it suitable for temporal prediction tasks such as disease progression. The results demonstrate that DyGraphTrans consistently outperforms state-of-the-art dynamic graph models across MCI to AD progression and high-acuity mortality prediction tasks. Beyond clinical EHR applications, DyGraphTrans generalizes well to benchmark dynamic graph datasets, achieving state-of-the-art performance on DBLP-3 and DBLP-10 while remaining competitive across diverse small and large-scale graphs. The results confirm that the proposed framework is robust while remaining memory-efficient on large-scale graph and capable of learning temporal dependencies effectively across both clinical data and non-EHR dynamic graph settings. Additionally, DyGraphTrans identifies the most influential time points and features for the prediction task. Although the current framework offers feature and time-point–level interpretability, extending it to patient-level explanations and inductive generalization to unseen patients remains an important direction for future work.

## Acknowledgements

This work was supported by the National Institute of General Medical Sciences of the National Institutes of Health under Award Number R35GM133657, the startup funds from the University of North Texas to S.B., summer Research Assistantships from the BioDiscovery Institute and Center for Computational Life Sciences to M.T.R. Data used in preparation of this article were obtained from the ADNI database (adni.loni.usc.edu). As such, the investigators within the ADNI contributed to the design and implementation of ADNI and/or provided data but did not participate in analysis or writing of this report. A complete listing of ADNI investigators can be found at: http://adni.loni.usc.edu/wp-content/uploads/how_to_apply/ADNI_Acknowledgement_List.pdf. The NACC database is funded by NIA/NIH Grant U24 AG072122, with data contributed by multiple NIA-funded Alzheimer’s Disease Research Centers (ADRCs). Full funding acknowledgments and center details are provided in the Supplemental Document.

## Data availability

The datasets were derived from sources in the public domain: ADNI: https://adni.loni.usc.edu/. NACC: https://naccdata.org/. MIMIC-IV: https://physionet.org.

## Author contributions

Conceptualization, M.T.R., S.B.; Methodology, M.T.R., S.B.; Data collection & processing, M.T.R., M.A.O.; Running experiments, M.T.R.; Writing-original Draft, M.T.R; Writing, Review & Editing, S.B., M.T.R.; Supervision: S.B.

