## Supplemental Document for "DyGraphTrans: A temporal graph representation learning framework for modeling disease progression from Electronic Health Records"

#### Supplementary Material

Most Tahmina Rahman
Mohammad Al Olaimat
Serdar Bozdag

Supplementary Table S1: SNF Hyperparameter values for EHR-based graph construction.

| Dataset | Network Type | Similarity Function | $K$ | $\mu$ | $t_{\text{diff}}$ |
|---|---|---|---|---|---|
| ADNI | Demographic | Jaccard | 8 | 1.0 | – |
| | MRI | Euclidean | 8 | 1.0 | – |
| | Cognitive | Euclidean | 8 | 1.0 | – |
| | Fused Network | – | 8 | – | 50 |
| NACC | Demographic | Jaccard | 10 | 0.8 | – |
| | Cognitive | Euclidean | 10 | 0.8 | – |
| | Fused Network | – | 10 | – | 50 |
| MIMIC-IV | Static | Jaccard | 5 | 0.5 | – |
| | Ventilation | Euclidean | 5 | 0.5 | – |
| | Labs & vital signs | Euclidean | 5 | 0.5 | – |
| | Fused Network | – | 5 | – | 20 |

### 1 Experiments

#### 1.1 Dynamic Graph Datasets

**EHR Datasets:** We evaluate DyGraphTrans on three longitudinal Electronic Health Record (EHR) datasets : ADNI, NACC, and the MIMIC-IV v2.2 intensive care cohort (Supplementary Table S2).

Supplementary Table S2: Dynamic homogeneous graph statistics for the ADNI, the NACC, and the MIMIC-IV datasets.

| Dataset | Patients (Nodes) | Static Features | Dynamic Features | Edges | Visits |
|---|---|---|---|---|---|
| ADNI | 137 | 6 | 20 | 6,573 | 7 |
| NACC | 440 | 7 | 5 | 232,320 | 8 |
| MIMIC-IV | 19,570 | 10 | 34 | 6,350,406 | 75 |

**Benchmark Dynamic Graph Datasets:** In addition to EHR data, we evaluate DyGraphTrans on widely used benchmark dynamic graph datasets spanning citation networks, social networks, and biological graphs (Supplementary Table S3). For each benchmark dataset, we constructed the final dynamic graph snapshots using the official GraphSSM [Li et al., 2024] preprocessing pipeline and strictly followed the same train/validation/test splits. The prediction task is temporal node classification, with supervision provided only at the final snapshot, consistent with the GraphSSM experimental protocol.

#### 1.2 Task Setup

**Task Setup for EHR Datasets:** For ADNI and NACC, models observe the first $t$ visits to predict the diagnosis at visit $t+1$. Specifically, for ADNI, the first six visits are used to predict the diagnosis at the seventh visit, while for NACC, the first seven visits are used to predict the diagnosis at the eighth visit. For MIMIC-IV, models use the first 48 hourly time points to predict in-hospital mortality at hour 75.

In all settings, temporal EHR graphs are constructed using patient attributes and their relationships, without incorporating outcome labels at intermediate visits. Supervision is provided exclusively by the label at the final time point, and evaluation is conducted on a held-out test set. All methods operate on sequences of fused dynamic graphs generated using Similarity Network Fusion (SNF). We adopt a 60–20–20 train/validation/test split, using identical data partitions across all models. Final results are reported as the mean and standard deviation over five independent runs using the same seed.

**Task Setup for Benchmark Temporal Graph Datasets:** For benchmark dynamic graph datasets, the task is temporal node classification. Models observe a sequence of historical graph snapshots and predict node labels at the final snapshot. Supervision is provided exclusively at the last time step, and no intermediate labels are used during training to prevent temporal label leakage. For each benchmark dataset, we strictly follow the same train/validation/test splits as GraphSSM to ensure a fair comparison.

Supplementary Table S3: Summary of benchmark dynamic graph datasets.

| Dataset | #Nodes | #Edges | #Features | #Classes | #Time Steps | Category |
|---|---|---|---|---|---|---|
| DBLP-3 [Xu et al., 2019] | 4,257 | 23,540 | 100 | 3 | 10 | Citation |
| Brain [Xu et al., 2019] | 5,000 | 1,955,488 | 20 | 10 | 12 | Biology |
| Reddit [Xu et al., 2019] | 8,291 | 264,050 | 20 | 4 | 10 | Society |
| DBLP-10 [Li et al., 2023] | 28,085 | 236,894 | 128 | 10 | 27 | Citation |
| arXiv [Hu et al., 2020] | 169,343 | 2,315,598 | 128 | 40 | 35 | Citation |
| Tmall [Li et al., 2023] | 577,314 | 4,807,545 | 128 | 5 | 186 | E-commerce |

##### 1.3 Baselines

**Baselines for EHR Datasets:** We compare DyGraphTrans with six state-of-the-art baselines.(1) EvolveGCN-H and (2) EvolveGCN-O use RNN to dynamically update the parameters of a GCN over time [Pareja et al., 2020]. Both architectures allow the GNN to evolve during inference.(3) GraphSSM is a state-space model for dynamic graphs that parameterizes temporal evolution using latent state transitions [Li et al., 2024].(4) WinGNN employs a sliding-window strategy to capture short-term temporal context while applying a GNN within each window [Zhu et al., 2023]. This design restricts memory growth and focuses the model on the most recent structural information.(5) ROLAND models graph evolution through an embedding update module that approximates temporal state transitions in GNN node embeddings and graph structure [You et al., 2022]. (6) TA-RNN is a deep learning model that operates on patient feature sequences [Al Olaimat et al., 2024]. TA-RNN introduces time-aware attention mechanisms within RNN to model irregular temporal intervals in EHR. All baselines are carefully tuned to achieve their best results. TA-RNN, EvolveGCN and GraphSSM implementations were taken from officially provided by the authors and the other baselines implementations were taken from Dynamic Graph Subtree Attention paper[Nguyen and Ta, 2025].

**Baselines for Benchmark Temporal Graph Datasets:** For the dynamic graph-based benchmark datasets, we additionally compare DyGraphTrans against a diverse set of graph methods. DeepWalk [Perozzi et al., 2014] and Node2Vec [Grover and Leskovec, 2016] are static graph embedding methods while HTNE [Zuo et al., 2018], M2DNE [Lu et al., 2019] and DynamicTriad [Zhou et al., 2018] are continuous-time temporal graph embedding approaches.MPNN [Panagopoulos et al., 2021], STAR [Xu et al., 2019], tNodeEmbed [Singer et al., 2019], EvolveGCN [Pareja et al., 2020], SpikeNet [Li et al., 2023] , ROLAND [You et al., 2022] and GraphSSM [Li et al., 2024] are discrete-time temporal graph based methods. For all benchmark experiments, the results of these methods were taken directly from the GraphSSM paper, and we reproduced our results following the same experimental setup, preprocessing, and evaluation protocol used in their study to ensure a fair comparison.

##### 1.4 Implementation Details

**Implementation Details for EHR Datasets:** DyGraphTrans is trained end-to-end using cross-entropy loss and the Adam optimizer, with early stopping based on validation performance. All baseline models are trained under identical preprocessing, graph construction, and evaluation procedures. We report Micro-F1 and Macro-F1 performance on held-out test sets, averaged over five independent runs with fixed random seeds.

**Implementation Details for Benchmark Temporal Graph Datasets:** For benchmark dynamic graph datasets, DyGraphTrans is trained using weighted

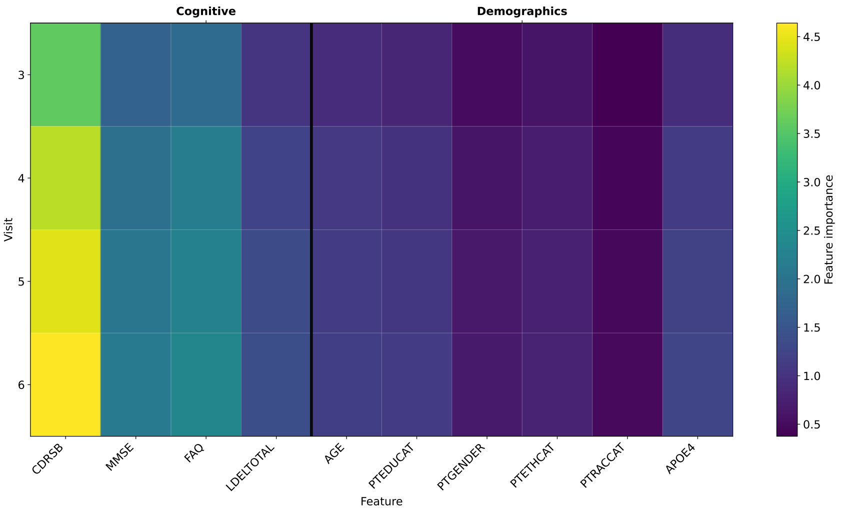


Supplementary Figure S1: Attention–gradient fused feature importance heatmap for NACC, aggregated over 20 random seeds (mean values). Each row corresponds to a time point (i.e., clinical visit), and each column corresponds to an EHR feature.

cross-entropy loss with the AdamW optimizer. Performance is evaluated using Micro-F1 and Macro-F1 scores on validation and test sets, and final results are reported as the mean and standard deviation over five independent runs with fixed random seeds.

#### 2 Acknowledgments for NACC data

The NACC database is funded by NIA/NIH Grant U24 AG072122. NACC data are contributed by the NIA-funded ADRCs: P30 AG062429 (PI James Brewer, MD, PhD), P30 AG066468 (PI Oscar Lopez, MD), P30 AG062421 (PI Bradley Hyman, MD, PhD), P30 AG066509 (PI Thomas Grabowski, MD), P30 AG066514 (PI Mary Sano, PhD), P30 AG066530 (PI Helena Chui, MD), P30 AG066507 (PI Marilyn Albert, PhD), P30 AG066444 (PI John Morris, MD), P30 AG066518 (PI Jeffrey Kaye, MD), P30 AG066512 (PI Thomas Wisniewski, MD), P30 AG066462 (PI Scott Small, MD), P30 AG072979 (PI David Wolk, MD), P30 AG072972 (PI Charles DeCarli, MD), P30 AG072976 (PI Andrew Saykin, PsyD), P30 AG072975 (PI David Bennett, MD), P30 AG072978 (PI Neil Kowall, MD), P30 AG072977 (PI Robert Vassar, PhD), P30 AG066519 (PI Frank aFerla, PhD), P30 AG062677 (PI Ronald Petersen, MD, PhD), P30 AG079280 (PI Eric Reiman, MD), P30 AG062422 (PI Gil Rabinovici, MD), P30 AG066511 (PI Allan Levey, MD, PhD), P30 AG072946 (PI Linda Van Eldik, PhD), P30 AG062715 (PI Sanjay Asthana, MD, FRCP), P30 AG072973

(PI Russell Swerdlow, MD), P30 AG066506 (PI Todd Golde, MD, PhD), P30 AG066508 (PI Stephen Strittmatter, MD, PhD), P30 AG066515 (PI Victor Henderson, MD, MS), P30 AG072947 (PI Suzanne Craft, PhD), P30 AG072931 (PI Henry Paulson, MD, PhD), P30 AG066546 (PI Sudha Seshadri, MD), P20 AG068024 (PI Erik Roberson, MD, PhD), P20 AG068053 (PI Justin Miller, PhD), P20 AG068077 (PI Gary Rosenberg, MD), P20 AG068082 (PI Angela Jefferson, PhD), P30 AG072958 (PI Heather Whitson, MD), P30 AG072959 (PI James Leverenz, MD).
